# Apparent Propagator Anisotropy from reduced Diffusion MRI acquisitions

**DOI:** 10.1101/798892

**Authors:** Santiago Aja-Fernández, Antonio Tristán-Vega, Derek Jones

**Affiliations:** Laboratorio de Procesado de Imagen (LPI), Universidad de Valladolid, Valladolid, Spain; Cardiff University Brain Research Imaging Center (CUBRIC), School of Psychology, University of Cardiff, UK

**Author notes:** Address correspondence to: Santiago Aja-Fernández, ETSI Telecomunicación, Campus Miguel Delibes s/n, Universidad de Valladolid, 47011 Valladolid, Spain. *Email address:* (Santiago Aja-Fernández).

**Keywords:** Diffusion MRI, propagator anisotropy, EAP, microstructure, HARDI

## Abstract

The Propagator Anisotropy (PA) is a measurement of the orientational variability inside a tissue estimated from diffusion MRI using the Ensemble Average diffusion Propagator (EAP). It is based on the quantification of the angular difference between the propagator in a specific voxel and its isotropic counterpart. The PA has shown the ability to reveal microstructural information of interest and meaningful descriptive maps inside the white matter. However, the use of PA is not generalized among the clinical community, due to the great amount of data needed for its calculation, together with the associated long processing times. In order to calculate the PA, the EAP must also be properly estimated. This task would require a dense sampling of the Cartesian q-space. Alternatively, more efficient techniques have been proposed in the last decade. Even so, all of them imply acquiring a large number of diffusion gradients with different b-values and long processing times.

In this work, we propose an alternative implementation to drastically reduce the number of samples needed, as well as boosting the estimation procedure. We avoid the calculation of the whole EAP by assuming that the diffusion anisotropy is roughly independent from the radial direction. With such an assumption, we achieve a closed-form expression for a measure similar to the PA but using information from one single shell: the Apparent Propagator Anisotropy (APA). The new measure remains compatible with standard acquisition protocols commonly used for HARDI (based on just one b-value). The intention of the APA is not to exactly replicate the PA but inferring microstructural information with comparable discrimination power as the PA but using a reduced amount of data.

We report extensive results showing that the proposed measures present a robust behavior in clinical studies and they are computationally efficient and robust when compared with PA and other anisotropy measures.

## 1. Introduction

The term Diffusion Magnetic Resonance Imaging (DMRI) refers to a set of diverse imaging techniques that, applied to brain studies, provide useful information about the organization and connectivity of the white matter. The most relevant feature of DMRI is its ability to measure orientational variance in the different tissues, i.e. anisotropy. Nowadays, the most common way to estimate the anisotropy is still via the popular diffusion tensor (DT) approach (Basser and Pierpaoli, 1996). DT-MRI brought to light one of the most common problems in DMRI techniques: in order to carry out clinical studies, the information given by the selected diffusion analysis method must be translated into some scalar measures that describe different features of the diffusion within every voxel. That way, metrics like the Fractional Anisotropy (FA) were defined with the DT as a starting point (Westin et al., 2002). Despite the strong limitations the underlying Gaussian assumption imposes to the diffusion model, the FA is still widely used in clinical studies involving DMRI.

Nevertheless, the diffusion mechanisms cannot be accurately described by DT-MRI because of the oversimplified Gaussian fitting. Accordingly, more evolved techniques with more degrees-of-freedom naturally arose, such as High Angular Resolution Diffusion Imaging (Tuch et al., 2003; Özarslan et al., 2006, HARDI) or Diffusion Kurtosis Imaging (Hansen and Jespersen, 2016, DKI). The trend over the last decade has consisted in acquiring a large number of diffusion-weighted images distributed over several shells (i.e. with several gradient strengths) and with moderate-to-high b-values to estimate more advanced diffusion descriptors, as the Ensemble Average Diffusion Propagator (Özarslan et al., 2013, EAP). The estimation relies more on model-free, non parametric approaches that can accurately describe most of the relevant phenomena associated to diffusion.

In order to estimate the EAP, a straight forward strategy like Diffusion Spectrum Imaging (Wedeen et al., 2005, DSI), would requires a huge number of images to attain a decent accuracy. Hence, alternative methods proposed from reduced (as opposed to dense) samplings of the q-space, being the most prominent Hybrid Diffusion Imaging (Wu and Alexander, 2007; Wu et al., 2008, HYDI), Multiple q-shell Diffusion Propagator Imaging: (Descoteaux et al., 2009, 2011, mq-DPI), Bessel Fourier Orientation Reconstruction (Hosseinbor et al., 2013, BFOR), the directional radial basis functions (Ning et al., 2015, RBFs), the Mean Apparent Propagator MRI (Özarslan et al., 2013; Avram et al., 2016, MAP-MRI) or the Laplacian-regularized MAP-MRI (Fick et al., 2016b, MAPL).

Regardless of the method selected to estimate the EAP, practical applications only use a reduce set of scalar measures derived from it: the probability of zero displacement, Q-space inverse variance, the return-to-plane and return-to-axis probabilities (Hosseinbor et al., 2012; Wu et al., 2008; Ning et al., 2015) or the Propagator Anisotropy (Özarslan et al., 2013, PA). In this work we will focus on the later.

Although the use of the PA is not generalized among the clinical community, there is a growing interest on the exploration of their potential clinical applicability, since it has shown the ability to reveal microstructural information of interest and meaningful descriptive maps of the white matter. Some seminal works have shown that the PA could be a valid biomarker for Alzheimer when evaluated over transgenic rats (Fick et al., 2016a). The same study also shows that the PA could be an important marker for longitudinal studies, since it uniformly changes over time, indicating a possible dependency with age. In Avram et al. (2016), PA shows higher tissue contrast than the FA and a more uniform behavior in white matter. Finally, the study carried out in Bernstein (2019) raises one of the problems of the PA: the bottleneck of studies with EAP-derived measures is the amount of data needed for the calculation. This issue, together with the long processing times needed for EAP estimation, has slowed down a widespread clinical adoption of such a measure.

The present paper delves into the hypothesis that the computation of the model-free EAP is not necessary to calculate the PA, on the contrary, a constrained model for radial diffusion may reveal analogous information using simpler protocols. To that end, we have first reformulated the inner product needed for the PA calculation for single-shell acquisitions based on different diffusion models, so that the corresponding scalar measures are compatible with more standard acquisition protocols like those used in HARDI or DKI, i.e., with faster acquisitions based on heavily reduced data sets and therefore applicable to data acquired within the clinical domain. Then, a novel diffusion anisotropy metric based on the PA is proposed, namely the Apparent Propagator Anisotropy (APA). Alternative implementations of the measure are also presented. The new metrics are extensively tested against PA and other anisotropy-based indices to check its capability to detect different configurations and its performance in the analysis of clinical data.

## 2. Theory

### 2.1. The Diffusion Signal

The EAP, *P*(**R**), is the Probability Density Function (PDF) of the water molecules inside a voxel moving an effective distance **R** in a time ∆. It is related to the normalized magnitude image provided by the MRI scanner, *E*(**q**), by the Fourier transform (Callaghan et al., 1988):

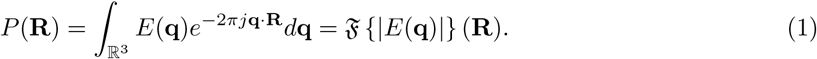

The inference of exact information on the **R**–space would require the sampling of the whole **q**–space to use the Fourier relationship between both spaces.

In order to obtain a closed-form analytical solution from a reduced number of acquired images, a model of the diffusion behavior must be adopted. The most common techniques rely on the assumption of a Gaussian diffusion profile and a steady state regime of the diffusion process that yields to the well-known Diffusion Tensor (DT) approach. Alternatively, a more general expression for *E*(**q**) can be used:

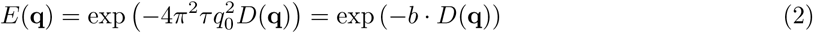

where the positive function *D*(**q**) = *D*(*q*_0_, *θ, φ*) is the Apparent Diffusion Coefficient (ADC), *b* = 4*π*^2^*τ*║**q**║^2^ is the so-called b-value and *q*_0_ = ║**q**║ and *θ, φ* are the angular coordinates in the spherical system. The effective diffusion time *τ* is defined as *τ* = ∆ − *δ*/3, where the diffusion time ∆ is usually corrected with the pulse duration *δ*. According to Basser (2002), in the mammalian brain, this mono-exponential model is predominant for values of b up to 2000 s/mm^2^ and it can be extended to higher values if appropriate multicompartment models of diffusion are used.

### 2.2. Inner Product and Propagator Anisotropy

Let *P*(**R**) and *Q*(**R**) be two different propagators. If we consider them as two different signals defined over a common signal space *S*, we can define an inner product as (Özarslan et al., 2013; Gallager, 2008)

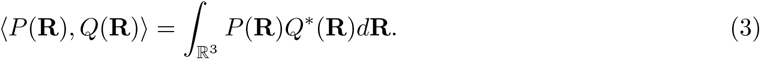

According to the Parseval Theorem (Gallager, 2008), since variables **R** and **q** are related via the Fourier Transform, there is an equivalence of this product in the **q**-space. Considering that the magnitude-reconstructed diffusion-weighted MR signal *E*(**q**) is always real and symmetric, *E^∗^*(**q**) = *E*(**q**) and *E*(**q**) = *E*(−**q**), we can write

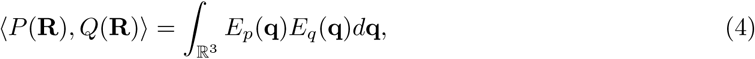

where 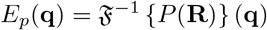 and 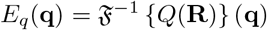. We can accordingly define the norm of a signal as

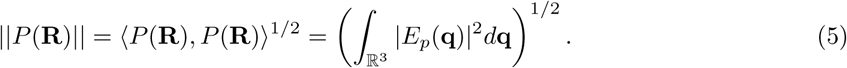

The *similitude* of two signals is given by the cosine of the angle between them, defined as

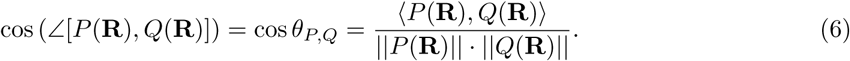

This measure can be used as an anisotropy measure using the EAP. In Özarslan et al. (2013), authors propose a measure called the Propagator Anisotropy (PA) which can be seen as a quantification of how a propagator diverges from the isotropic one. It is defined as

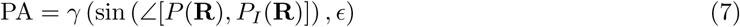

where *P*(**R**) is the actual propagator and *P*_*I*_(**R**) and equivalent isotropic propagator. The function *γ* (*., ϵ*) is a nonlinear transformation to better distribute the output values in the range [0, 1]. The sine is calculated from the cosine in eq. (6) as:

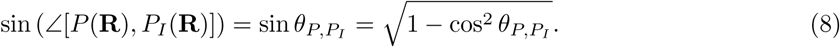

### 2.3. Apparent Propagator Anisotropy

As previously stated, one of the problems in the calculation of the PA is the large amount of measures needed in order to accurately estimate the EAP from the samples. In order to use a limited amount of acquisitions to estimate a similar anisotropy metric, we assume a prior model that assures that it can be carried out using data collected over one single shell. To that end, we are forced to consider that the diffusion *D*(**q**) does not depend on the radial direction, i.e. *D*(**q**) = *D*(*θ, φ*), so that eq. (2) becomes:

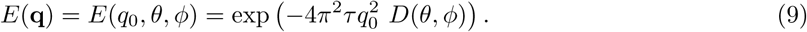

Note that, although *D*(**q**) no longer depends on *q*_0_, *E*(**q**) does. This assumption, although restrictive, is used to define certain diffusion modalities in HARDI (Descoteaux et al., 2006; Özarslan et al., 2006), where only one shell is usually acquired. This simplification was initially intended to overcome the limitations of the DT by allowing the diffusion to be evaluated across many orientations, as opposed to the single orientation described by the DT.

In what follows, we redefine the inner product using the simplification in eq. (9) and we use it to define an anisotropy metric related to the PA for a specific shell.

**First**, we define the isotropic equivalence of the signal, *E*_*I*_(**q**) as

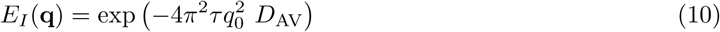

where *D*_AV_ is the *average diffusivity*. It can be seen as the as the value of the ADC over a unitary sphere:

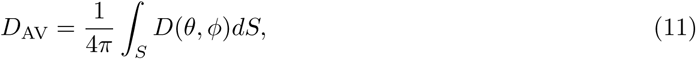

In order to calculate the integration on the surface of the sphere from a limited number of samples we use a Spherical Harmonics (SH) decomposition:

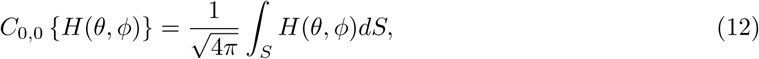

and therefore the *D*_AV_ can be calculated as:

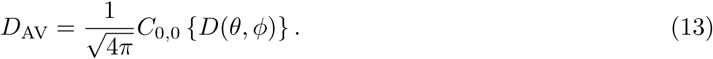

**Second**, we calculate the norm of *P*(**R**) and *P*_*I*_(**R**) under the considered assumption:

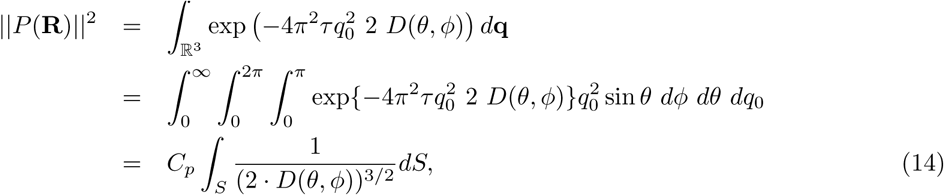

where *C*_*p*_ is a constant. Using a SH decomposition we can write

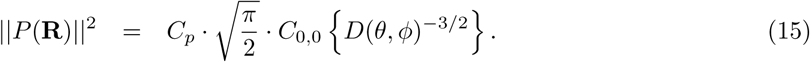

Following the same reasoning, the norm of the isotropic equivalent is:

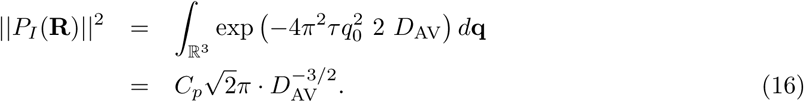

**Third**, we calculate the inner product of both signals using the single shell assumption:

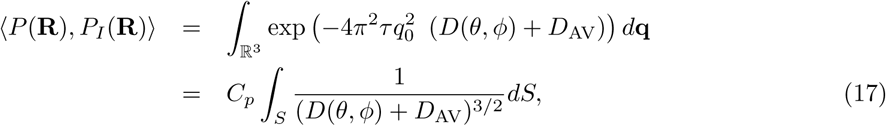

And using the SH decomposition for the calculation of the integral:

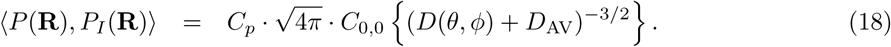

Next, we calculate the cosine and sin of the angle between both signals

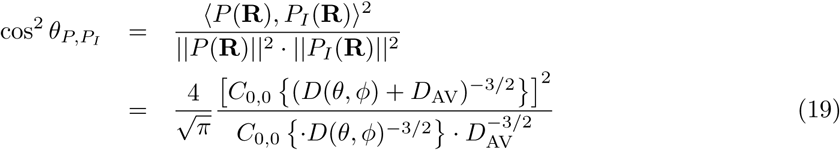

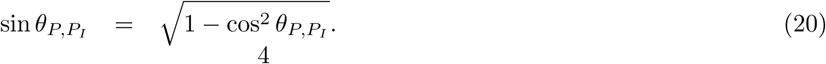

**Finally**, the PA is calculated using the Gamma transformation proposed by Özarslan et al. (2013):

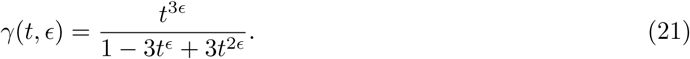

Since the measure is grounded on a initial assumption, the value could vary for different shells. This way, the Apparent Propagator Anisotropy (APA) at a given b-value is calculated as

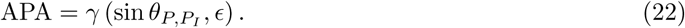

We define also the measure without the non-linear transformation as:

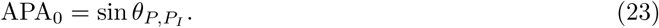

### 2.4. Dependence with Acquisition Parameters

The measure previously defined can be subject to acquisition artifacts, and it will be corrupted with acquisition noise. The presence of noise will introduce a bias in the estimator that depends on the signal-to-noise ratio (SNR) and may also depend on other acquisition parameters. In what follows we will analyze the dependence of the bias with those acquisition parameters and, in particular, its dependence with the b-value.

The signal *E*(**x**) is the acquired magnitude signal *S*_*i*_(**x**) normalized by the baseline *S*_0_(**x**):

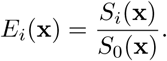

For the sake of simplicity, we assume that the acquired signals *S*_*i*_(**x**) are corrupted with Rician noise (Aja-Fernández and Vegas-Sánchez-Ferrero, 2016):

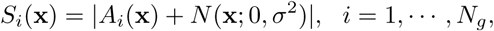

where *A*_*i*_(**x**) is the original signal if no noise is present and *N*(**x**; 0, *σ*^2^) is a complex additive Gaussian noise with zero mean and variance *σ*^2^. This is a common assumption in MRI acquisitions, valid for single–coil acquisitions and multi–coil parallel imaging reconstructed with a spatial matched filter, like SENSE, for instance. In the latter, noise can become *non–stationary*, i.e., the variance of noise will depend on the position and *σ* must be replaced by *σ*(**x**), which does not affect to the following study. We can also assume that the SNR in the baseline is high enough so we can consider *S*_0_(**x**) a *noiseless image*.

For the sake of simplicity in the analysis, we will estimate the bias of the cos^2^ in eq. (19) instead of APA, since it is easier to calculate and the bias will be related. The analytical study of the bias is described in Appendix A.

To better understand the effect of the SNR and the b-value, let us simplify the resulting bias by assuming an isotropic diffusion. This way, we can write the mean of the measure as

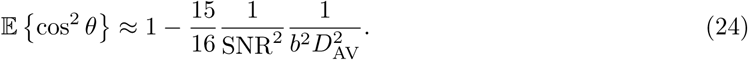

Therefore, the estimation bias of the cos^2^ *θ* can be quantified, for the isotropic case as:

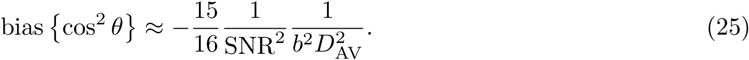

Note that the SNR is the signal to noise ratio at the acquired DWI. We can write it as

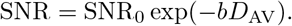

where SNR_0_ is the SNR measured at the baseline. So, we can rewrite the bias as:

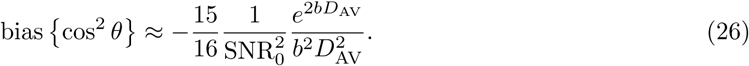

Although this is a very restricted case, it gives us a very helpful insight of the dependence of the measure with b and SNR. In fig. 1 the bias in eq. (26) is depicted as a function of the SNR (for a fixed diffusivity) and as a function of the ADC (for a fixed SNR=10).

**Figure 1:**
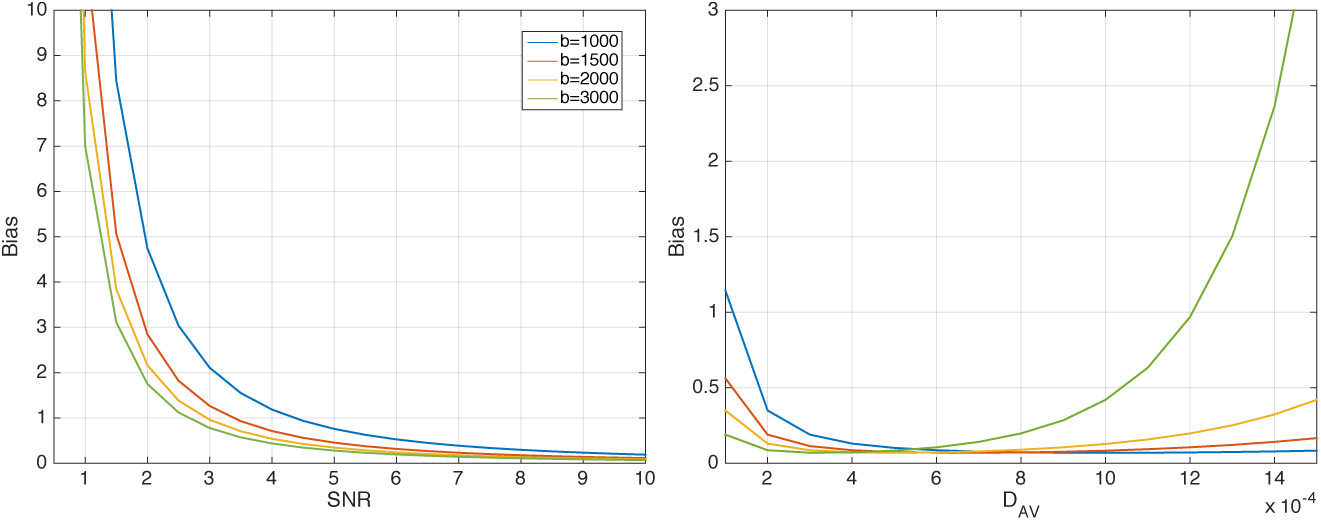
Bias of cos^2^ *θ* for the isotropic case as a function on the b-value and the SNR. (a) *D*_AV_ = 3 × 10^−4^ (b) SNR=10

Note that, for the same diffusion, there is a different bias for different values of b. For low SNR, this bias can grow, but when the SNR grows, the values for the different b converge. However, as we can see in the fig. 1-(b), the differences between values are also a function of the ADC: different values of the ADC can produce different bias for different b values. We will test this on real data in the experiments section.

### 2.5. Relation of APA with other anisotropy measures

The APA can be seen as the projection of diffusion *E*(**q**) over an isotropic equivalent considering just one shell and no radial diffusion. The same idea can be extrapolated to the diffusivity *D*(*θ, φ*). Thus, the equations can be modified accordingly:

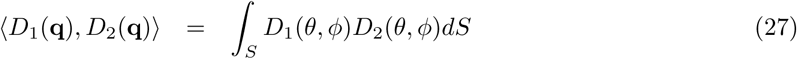

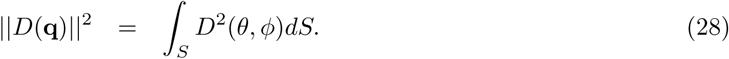

If we do not consider the Gamma transformation, the Diffusion Anisotropy (DiA) can be defined as

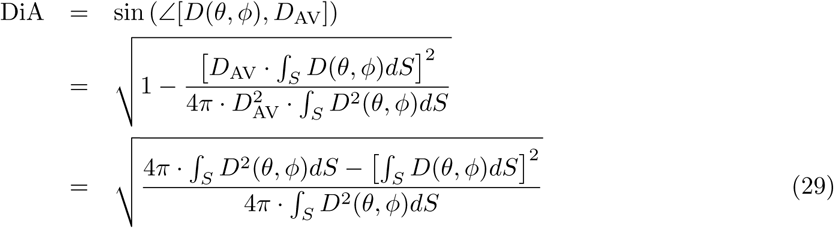

Note that the dependency with *D*_AV_ disappears. Using the SH decomposition for the calculation of the integral, we can write:

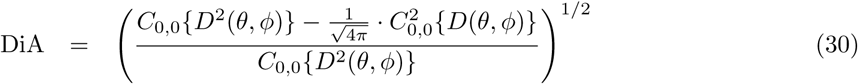

The DiA so defined, can be seen as a generalization of the Coefficient of Variation of the Diffusion (CVD) defined in Aja-Fernández et al. (2018) as a robust alternative for the FA. According to Tristán-Vega (2009), this measure is as an alternative implementation of the Generalized Anisotropy (Özarslan et al., 2005):

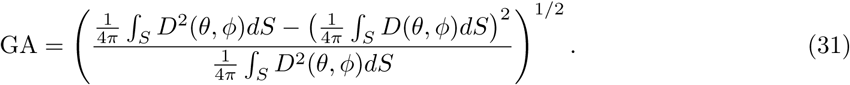

In order to increase the dynamic range of the metric, we can also use the *γ*(*t, ϵ*) transformation in eq. (21). This way, the DiA can be alternatively defined as:

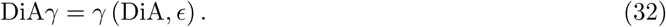

An overview of all the proposed diffusion anisotropy metrics, together with the specific numerical implementation, is presented in Table 1.

**Table 1:** Survey of the proposed anisotropic diffusion metrics.

| Measure | Formula | Practical implementation |
| --- | --- | --- |
| $D_{AV}$ | $\frac{1}{4\pi} \int_S D(\theta, \phi) dS$ | $\frac{1}{\sqrt{4\pi}} C_{0,0} \{D(\theta, \phi)\}$ |
| $APA_0$ | $\sqrt{1 - \frac{[\int_S (D(\theta, \phi) + D_{AV})^{-3/2} dS]^2}{\sqrt{2\pi} D_{AV}^{-3/2} \int_S (2D(\theta, \phi))^{-3/2} dS}}$ | $\sqrt{1 - \frac{4}{\sqrt{\pi}} \frac{[C_{0,0} \{(D(\theta, \phi) + D_{AV})^{-3/2}\}]^2}{C_{0,0} \{D(\theta, \phi)^{-3/2}\} \cdot D_{AV}^{-3/2}}}$ |
| APA | $\gamma(APA_0, \epsilon)$ | $\gamma(APA_0, \epsilon)$ |
| DiA | $\sqrt{\frac{4\pi \cdot \int_S D^2(\theta, \phi) dS - [\int_S D(\theta, \phi) dS]^2}{4\pi \cdot \int_S D^2(\theta, \phi) dS}}$ | $\sqrt{1 - \frac{\frac{1}{\sqrt{4\pi}} \cdot C_{0,0}^2 \{D(\theta, \phi)\}}{C_{0,0} \{D^2(\theta, \phi)\}}}$ |
| DiAγ | $\gamma(DiA, \epsilon)$ | $\gamma(DiA, \epsilon)$ |

## 3. Experiments and Results

### 3.1 Setting-up of the experiments

For the following experiments, the different methods are implemented using SH expansions of even orders up to 6 in all cases (when needed), with a Tikhonov regularization parameter *λ* = 0.006. PA is calculated using the DIPY toolbox with anisotropic basis and radial order 6 Özarslan et al. (2013).

Four different real data sets are used for the experiments:

- Human Connectome Project (HCP)^1^ are considered, specifically, five different volumes: MGH 1007, MGH 1010, MGH 1016, MGH 1018 and MGH 1019, acquired in a Siemens 3T Connectome scanner with 4 different shells at *b* = [1000, 3000, 5000, 10000] s/mm^2^, with [64, 64, 128, 256] gradient directions each, in-plane resolution 1.5 mm and slice thickness was 1.5 mm. The acquisition included 40 different baselines that were averaged to reduce the noise^2^.
- Public Parkinson’s disease database (PDD): public available data base^3^ acquired in the Cyclotron Research Centre, University of Liège. It consists on 53 subjects in a cross-sectional Parkinson’s disease (PD) study: 27 PD patients and 26 age, sex, and education-matched control subjects. Data were acquired on a 3 T head-only MR scanner (Magnetom Allegra, Siemens Medical Solutions, Erlangen, Germany) operated with an 8-channel head coil. Diffusion-weighted (DW) images were acquired with a twice-refocused spin-echo sequence with EPI readout at two distinct b-values (b = 1000, b = 2500 s/mm^2^) along 120 encoding gradients that were uniformly distributed in space by an electrostatic repulsion approach. For the purposes of motion correction, 22 unweighted (b = 0) volumes, interleaved with the DW images, were acquired. Acquisition parameters are TR=6800 ms, TE=91 ms, and FOV=211 mm^2^, no parallel imaging and 6/8 partial Fourier were used. More information can be found in Ziegler et al. (2014).
- Multi–parameter dataset (GMH): acquired in a PHILIPS 1.5 T MR scanner at the Gregorio Marañón Hospital (Madrid, Spain). Thirteen healthy male adults, aged between 23 and 31 (average age 27 years), participated in this study. DWIs were acquired using a multi-shot pseudo-3D double spin-echo echo-planar imaging (SE-EPI) sequence. Each exam was composed of different DTI acquisitions with different combination of parameters. Two different b-values were used: 800, and 1300 s/mm2. All the scans were acquired with 61 gradient directions and one baseline volume. The gradient directions were specifically acquired so that they can be subsampled to 40, 21 or 6 gradient directions while remaining equally spaced for each configuration. This subsampling technique allows the measurement of the effect of different number of gradients with only one data acquisition. The scans were acquired with a spatial resolution of 2.5 2.5 2.5 mm^3^. Other diffusion acquisition parameters are: echo time (TE) 1.6 ms, repetition time (TR) 8 ms. More information can be found in Barrio-Arranz et al. (2015).
- Multishell data acquired at CUBRIC (CBR): 14 healthy volunteers scanned in a 3T Siemens Prisma scanner (80 mT/m) with a pulsed-gradient spin-echo (PGSE) sequence. Three shells were acquired at b=[1200, 3000, 5000] s/mm^2^ with 60 directions per value. The resolution is isotropic of 1.5 mm^3^. Other acquisition parameters are: TE=80 ms, TR=4500ms, ∆/*δ* = 38.3/19.5 ms, parallel imaging acquisition (GRAPPA2) with sum of squares combination and 32 channels.

### 3.2. Visual Results

First, a visual comparison of the metrics is done using 3 slices (42,52,65) from the HCP volume (MGH1007). The proposed measures (APA, APA_0_,DiA and DiA*γ*) are calculated using a single shell for b=3000 s/mm^2^. For the sake of comparison, we have also calculated FA at b=1000 s/mm^2^, GA at b=3000 s/mm^2^ and PA using all the available information (4 shells). Results are shown in Fig. 2. All the metrics show a similar look, highlighting those anisotropic areas inside of the white matter. APA_0_ and DiA, as expected, show little contrast, a fact that is corrected by APA and DiA*γ*. However, it is not the visual aspect what we are interested in, but the ability to discriminate differences inside the white matter.

**Figure 2:**
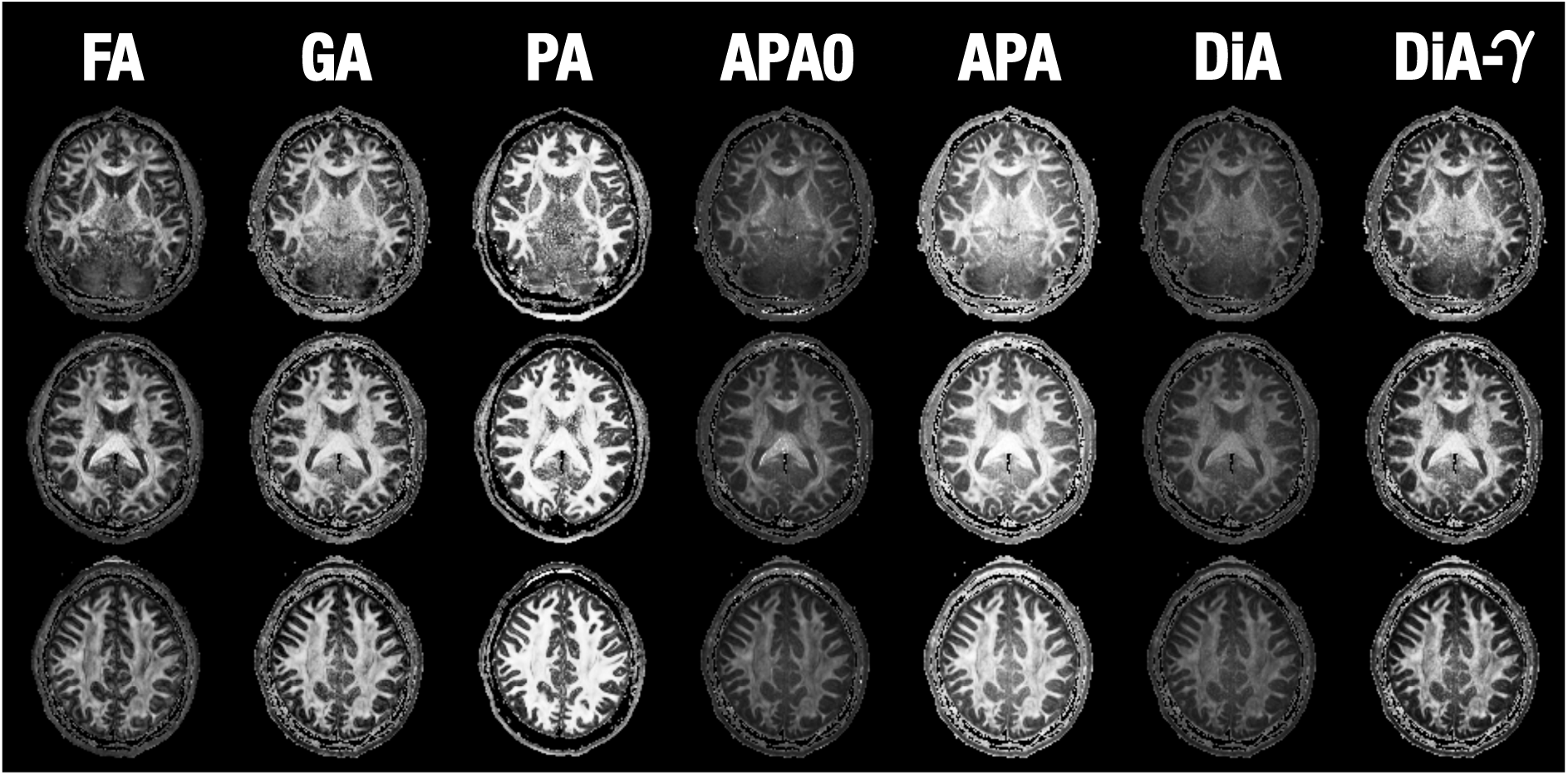
Visual comparison of the diffusion anisotropy metrics using slices 42, 52 and 65 of of the MGH1007 volume from HCP. FA is calculated using b=1000 s/mm^2^, GA, PA, SIN, DiA and DiA*γ* using b=3000 s/mm^2^ and PA using 4 shells (1000, 3000, 5000 and 10000 s/mm^2^).

### 3.3. Validation with Clinical Data

Next, we intend to test the clinical potential of the new metrics, for which we have explored the PPD database. Parkinson disease is known to affect the substantia nigra or the gray matter more than white matter. However, significant differences have also been reported in several white matter regions such as the corpus callosum (CC), the corticospinal tract, or the fornix Atkinson-Clement et al. (2017). Since the aim of this experiment is testing the capability of the proposed metrics to probe the micro-structural properties of the white matter, we have accordingly focused on commonly-studied white matter tracts.

FA is calculated as a reference value using MRTRIX^4^ with the data at b=1000 s/mm^2^. The FA maps of all the volumes are warped to a common template using the standard TBSS pipeline Smith et al. (2006). The same transformation is applied to all the metrics considered for the experiment (APA, APA_0_, DiA, DiAγ, PA and GA). Two different analysis are considered:

1. A voxelwise cross-subject analysis using the FA skeleton with the randomise tool from the FSL toolbox (which performs a nonparametric permutation inference over the data) with 500 realizations. Those voxels with p*<* 0.01 are highlighted in Fig. 3. In red, those points where the considered metric decreases in the PD with respect to the controls.

**Figure 3:**
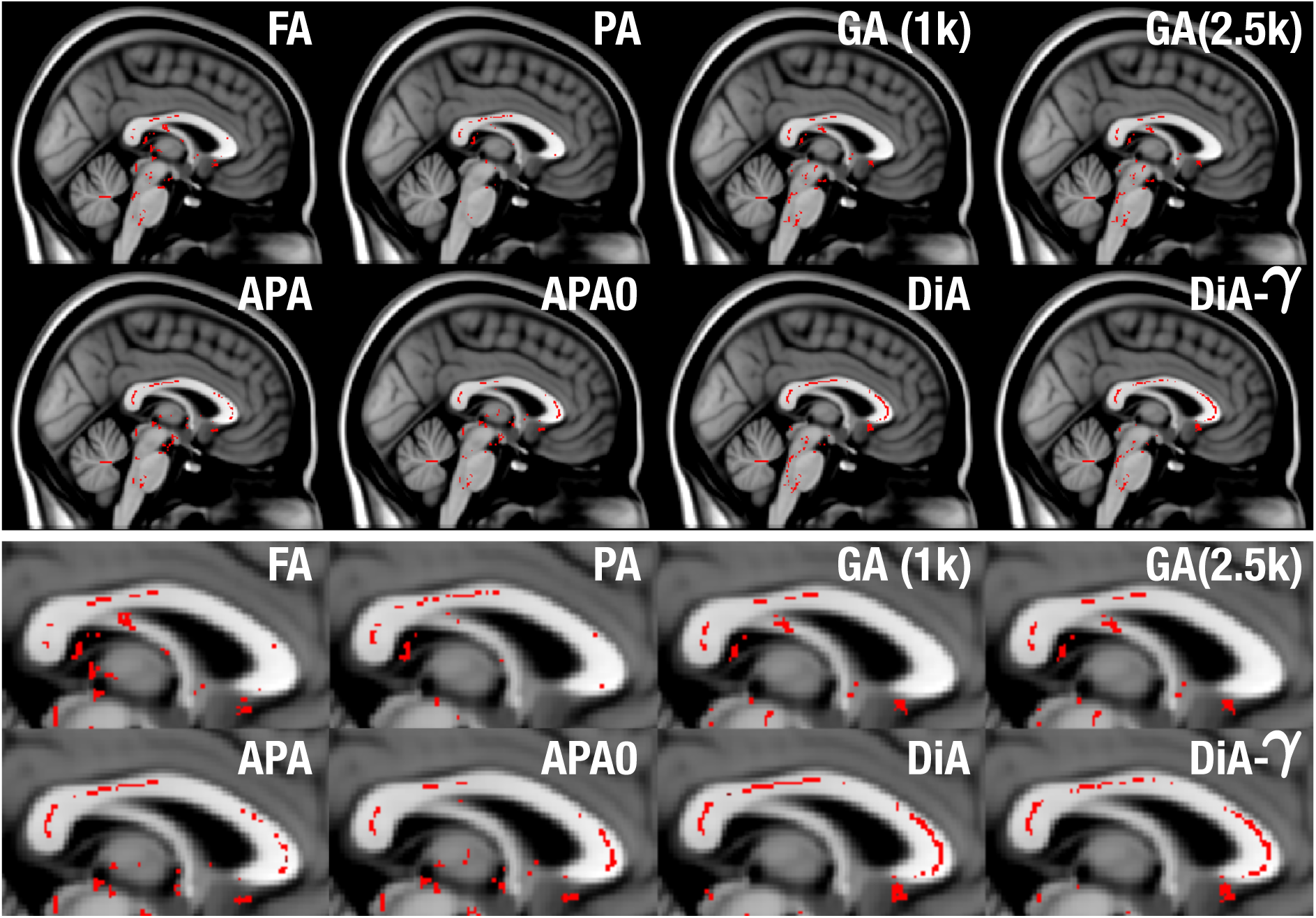
Significant differences found by statistical test for the Parkinson database, using a voxelwise analysis over the FA skeleton for the different considered metrics (sagittal view). In red, those points where the considered metric decreases in the PD with respect to the controls with statistical significance above 99% (p< 0.01).
2. A region of interest oriented analysis. The three regions of the CC (genu –GCC–, body –BCC–, and splenium –SCC–) are identified on the subjects using the JHU WM atlas Mori et al. (2005). The average value of the different measures inside each ROI is calculated using the values similar to the 2% and 98% percentiles. First, effect sizes were estimated using the Cohen’s d. Results are depicted in Fig 4. Then we carry out a two-sample, pooled variance *t*-tests between controls and patients for each of the measures considered and at each of the three sections of the CC segmented in the JHU WM. Table 2 shows the results.

**Table 2:** Two-sample, pooled variance, *t*-tests for each measure and at each section of the corpus callosum: GCC (genu), BCC (body), and SCC (splenium). The *p*-values represent the probability that the averaged values (using the values between the 2% and 98% percentiles) of each region of the corresponding tract have identical means for both controls and patients. Differences with statistical significance above 99% are highlighted in green, and those with significance over 95% are highlighted in amber.

|  | B val | GCC | BCC | SCC |
| --- | --- | --- | --- | --- |
| FA | 1000 | 0.378 | 0.205 | 0.192 |
| MAPMRI-PA | all | 0.656 | 0.585 | 0.517 |
| MAP-PA-DTI | all | 0.664 | 0.290 | 0.345 |
| GA | 1000 | 0.443 | 0.151 | 0.102 |
|  | 2500 | 0.063 | 0.078 | 0.015 |
| APA | 1000 | 0.555 | 0.296 | 0.310 |
|  | 2500 | 0.038 | 0.238 | 0.116 |
| APA<sub>0</sub> | 1000 | 0.309 | 0.676 | 0.436 |
|  | 2500 | 0.180 | 0.472 | 0.062 |
| DIA-γ | 1000 | 0.448 | 0.125 | 0.110 |
|  | 2500 | 0.093 | 0.099 | 0.029 |
| DIA | 1000 | 0.431 | 0.183 | 0.047 |
|  | 2500 | 0.057 | 0.071 | 0.004 |

**Figure 4:**
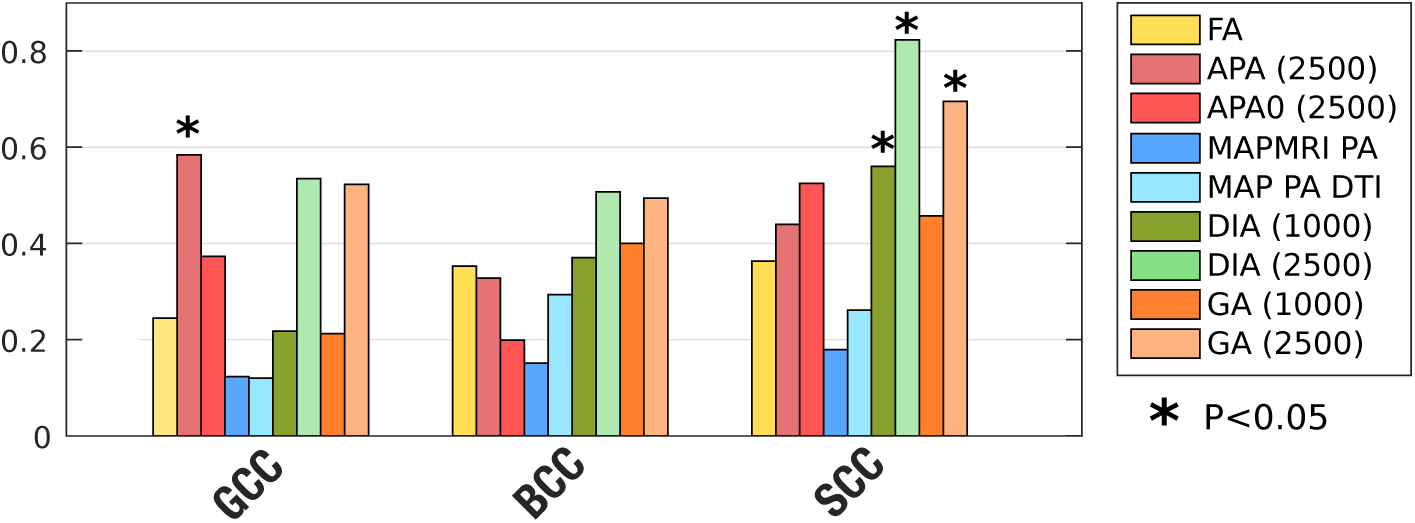
Absolute value of effect sizes (Absolute Cohen’s d) for associations between PD and controls in the Parkinson data base.

Since the aim of this experiment is testing the capability of the proposed measures to probe the micro-structural properties of the white matter, we have accordingly focused on the CC, where previous studies have reported main differences between PD and healthy controls. If we focus on this area in a sagittal plane, see Fig. 3, we can see that the FA and GA only find some isolated voxels with statistically significant differences. PA finds some extra voxels, but it cannot show its true potential due to the small *b*-values considered (higher *b*-values should result in more accurate EAP estimates). In contrast, the proposed measures show more differences along the whole CC. All of them, specially DiA and DiA*γ* find differences in the Genu of the CC (GCC).

In the region-of-interest analysis, it is precisely in the same area SCC that all the measures show the greatest values of Cohen’s d, see Fig. 4. Once again, DiA is the ones showing larger effect sizes, although in this area, GA (with b=2500) is also able to find significant differences, see Table 2. However, note that DiA shows a statistical significance above 99%. If we focus on the GCC, in this ROI analysis, only APA is able to find differences. On the other hand, note that PA calculated with MAPMRI and the DTI version show very low effect sizes and they are not able to detect significant differences in any part of the CC. This is related to the need of the measures of great amounts of information to properly estimate the EAP and then derive the PA. The use of only two shells, although possible, is not enough for an accurate estimation. Hence, the need of alternative measures for these scenarios.

### 3.4. Sensitivity analysis to acquisition parameters

The initial assumption for APA is that *D*(*θ, φ*) does not heavily depend on the b-value. However, we have shown that the bias on the measure depends on the SNR and on the b-value. So, next, we carry out three different experiments to quantify the sensibility of the proposed measures to the b-value.

First, in section 2.4 and appendix A the bias of the cos^2^ *θ* is quantified for high SNR. In order to test the model, a simulation is done: two voxels with isotropic and three anisotropic ones are simulated, using three different b values: 1000, 1500 and 2000 s/mm^2^. For the isotropic voxels two diffusivity values are considered, *D*_ISO_ = 300*µ*m^2^/s and *D*_ISO_ = 1000*µ*m^2^/s. The anisotropic ones are created with a tensor model with the greatest eigenvalue of value *D* and radial eigenvalues of value *d*, see fig 5.

**Figure 5:**
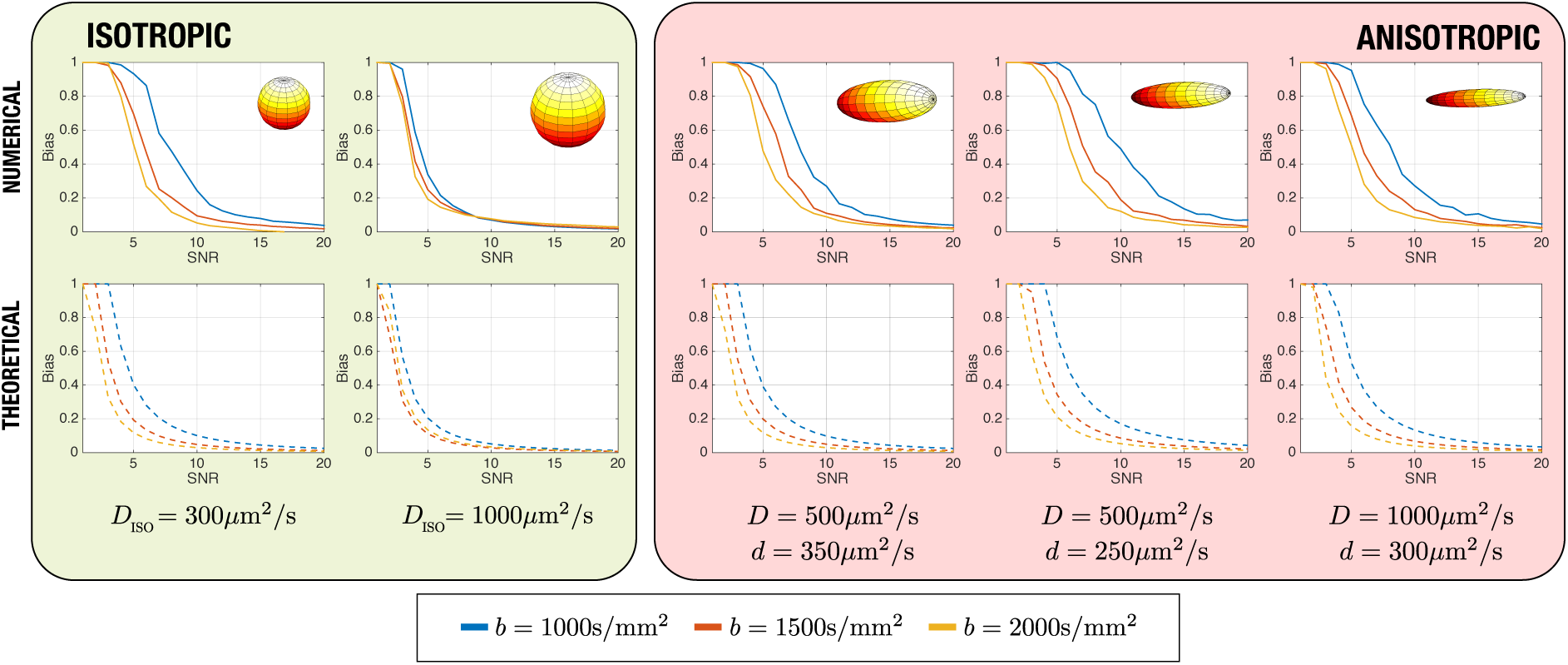
Bias of the estimation of cos^2^ *θ* for 5 different cases.

For the experiment, the bias for different SNR values is calculated for the simulations, together with the theoretical value. Results are depicted in Fig. 5. Note that the bias decreases in all cases when the SNR increases. However, in the anisotropic case, the decrease is slower; the differences of bias for different b values can be see for higher SNR. In addition, note that although the theoretical values are not exactly the numerical ones, the trend and the separation for different b values can still be perceived.

Some conclusions may be raised from these results: (1), the bias and the dependence with b decreases with the SNR. If the SNR is high enough, the bias will disappear. (2) The bias is smaller for higher b values. However, note that the SNR also depends on the b value. So, a reduction of the bias cannot be achieved by only increasing b.

These preliminary results give an interesting insight of what is happening with the measures. However, we will now move to real data, to better quantify the effect of changing the b value. Data from the CBR database is used for the next experiment. Each volume is divided in 6 different regions according to their diffusion features. To do that, the APA is first calculated and those voxels with APA< 0.1 are removed. The remaining voxels are clustered in 6 different groups using k-means, being the resulting centroids: *C*_*L*_ = 0.21, 0.35, 0.48, 0.63, 0.78, 0.93. Each voxel in the white matter is assigned to one cluster using its PA value and the minimum distance. All the proposed anisotropic diffusion measures are computed for each shell, and the median value inside each of the five clusters is depicted in Fig. 6.

**Figure 6:**
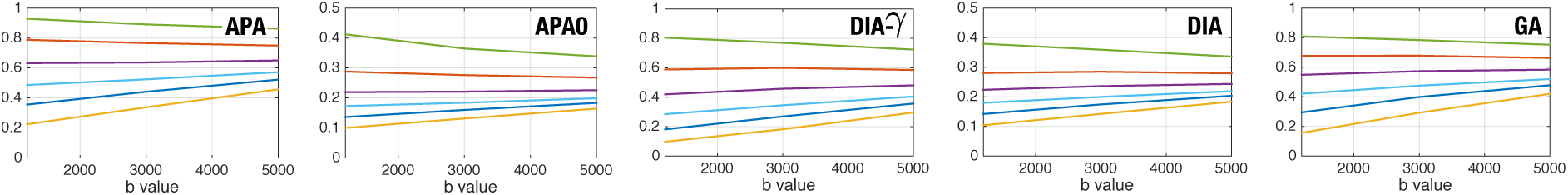
Evolution of the proposed measures with the b-value, using data from the A 3T PRISMA SCANNER. The volume has been clustered in 6 different sets and the median of each set is shown. Centroids of the data *C*_*L*_ = {0.21, 0.35, 0.48, 0.63, 0.78, 0.93}.

All of the measures show a dependence with the b-value: the smallest values values increase together with the b-value, whereas the higher values show a decrease. However, and this is the key point, the separation between clusters remains for different b values. This means that the differences in the anisotropy detected by these measures can be detected when using different shells.

Finally, we aim to quantify the effect of the variation of the number of gradients together with the b-value on real data. To that end, we consider the GMH data set, acquired with different acquisition parameters. The FA maps of all the volumes are warped to a common template using the standard TBSS pipeline. 48 different white matter regions of interest are identified on the subjects using the JHU WM atlas Mori et al. (2005). Different anisotropy measures are calculated for each region, with different b values and different number of gradients. The average value of the different measures inside each ROI is calculated using the values similar to the 5% and 95% percentiles. Then we carry out a two-sample, pooled variance *t*-tests between each metric calculated with different acquisition parameters for each of the regions. This way, we can quantify if the same volumes are detected as different when using different acquisition parameters. Table 3 shows the results.

**Table 3:**
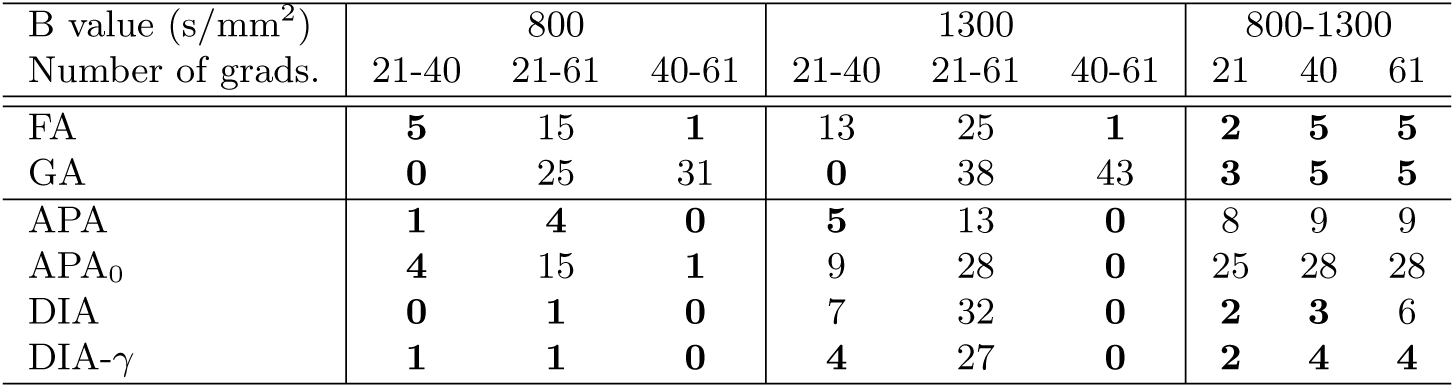
Number of ROIS considered to be different with statistical significance above 99% (out of the 48 regions defined in the JHU WM atlas) for the dataset GMH for different acquisition parameters.

From the results we can see that, as expected, when the number of acquired directions highly differ (21 vs 61 gradients) a great number of differences arises for all the metrics, specially for the higher b value, b=1300 s/mm^2^. For the same b value, a smaller variation in the directions (40 vs 61) gives much better results is most metrics, except the GA. The proposed metrics seem to be robust to this small changes. When the change is between 21 and 40, the numbers drop again, since 21 gradient did to capture all the directional information needed for the accurate estimation of APA and DiA. On the other hand, as expected from the previous experiment, APA and APA0 are very sensitive to the changes in the b-value. DiA shows here a most robust behavior.

### 3.5. Execution Times

The long processing times associated to the estimation of EAP-based measures is one of the issues that has slowed down a widespread clinical adoption of the PA. Precisely, the linear nature of SH needed to estimate the APA usually yields to a significant reduction of the calculation time, that can be several orders of magnitude faster than whole EAP-based techniques.

To test this extreme, a volume from the PDD is used here to compute APA and PA measures on a quad-core Intel(R) Core(TM) i7-4770K 3.50GHz processor under Ubuntu Linux 16.04 SO. PA is calculated using the two available shells with MAP-MRI using the DIPY library under Python 3.6.4 (scipy 1.0.0)^5^. APA is implemented using one single shell in MATLAB without multi-threading. The results are reported in Table 4.

**Table 4:** Estimated execution times for the calculation of APA and PA for a single volume.

| Method | Time |
| --- | --- |
| PA (MAP-MRI) | 2h 53min |
| APA | **3.17s** |

Though raw execution times are an ambiguous performance index (they can be dramatically improved, for example, via GPU acceleration), they give a reasonable idea of the complexity of each method. The calculation of the APA for the whole volume is almost instantaneous, which makes it feasible for practical studies.

## 4. Discussion and conclusions

The intention of the new anisotropy measure proposed, APA, as well as the other related measures derived, is not to exactly replicate a measure like the PA but, using a similar philosophy, inferring microstructural information with comparable discrimination power as the PA estimated using EAP-based methods. The original PA calculated from the EAP explicitly account for the radial behavior of the diffusion signal, which is actually sampled. For the APA calculation, the radial behavior is not sampled but modeled as a mono-exponential decay. Initially we can think that the computation of the whole EAP would provide a more specific and sensitive measure, since the anisotropy information encoded in the radial direction would be neglected in APA. Although this could be the case for a dense sampling of the **q** space, results for the clinical data (see the PDD experiment) paradoxically show the opposite. When only two shells are available, i.e. there is a reduced acquired data set, there is not enough information to accurately estimate the EAP and, consequently, a smooth version of the actual EAP is estimated instead. As a result, in these environments, the PA has shown a power to resolve microstructural features even below the capabilities of conventional DT-MRI.

On the other hand, experiments carried out in this paper confirm that the proposed measures show a discriminant power over traditional DT markers and, in some occasions, even over the PA. The aim of the experiment is not demonstrating the clinical usefulness of the new measures in the particular case of PD, but testing its capability to describe microstructural features when compare with state-of-the-art methods. We are aware that the finding of more significant differences does not directly implies that one method is better than other. But we can assure that the new measures are more sensitive to changes between groups. It will require further clinical studies to determine how this differences are linked to structural differences.

The main advantage of the proposed measures, when compared to PA, is that they can be calculated from a reduced set of measures. Initially they are intended for one shell, but the methodology can be easily extrapolated to more that one. In addition, the experiments with the GMH data set have shown a great robustness to the variability on the number of gradient directions, specifically between 40 and 61, which will allow a further reduction of the acquired data, compatible with nowadays standard acquisition protocols, with as few as 64 gradient directions. It is a common practice acquiring two shells (b=[1000, 3000] s/mm^2^, for instance) to estimate classical diffusion parameters, like the FA and MD, but also advanced models (DKI, HARDI, CHARMED, etcetera). APA and the additional anisotropy measures proposed can also be calculated with no additional effort and without changing the acquisition protocol.

Moreover, since the computation of APA avoids the estimation of the actual EAP it can be done in a fast and robust way, i.e., without imposing a computational burden to the standard protocols, see Table 4. A whole volume can be processed in a matter of seconds while the processing of the original PA usually take some hours, which obviously limits its applicability.

On the other hand, the major drawback of APA is the explicit assumption of a specific radial behavior for the diffusion, which cannot fit the whole q-space. As a consequence, the selection of the b-value may change the values of the measures. However, as we have shown, although the values change with b, the differences between areas remain. This implies the results of clinical trials could be compared against each other only if the same b-value is preserved across the studies. This is by no means something new to diffusion imaging: it is well-known that a change in the acquisition parameters (number of gradients, b-value, resolution, scanner vendor, etcetera) seriously affects scalar measures like the FA or the MD (Aja-Fernández et al., 2018; Barrio-Arranz et al., 2015).

## Acknowledgments

This work was supported by Ministerio de Ciencia e Innovación of Spain with research grants RTI2018-094569-B-I00 and PRX18/00253 (Estancias de profesores e investigadores senior en centros extranjeros). The authors thank the contributors of DIPY project (http://nipy.org/dipy/) for providing the MAP-MRI basis implementation and specially to Rutger Fick for his implementation for PA calculation and interesting discussion about MAP-MRI model.

Data collection and sharing for this project was provided by (1) the *Human Connectome Project* (HCP; Principal Investigators: Bruce Rosen, M.D., Ph.D., Arthur W. Toga, Ph.D., Van J. Weeden, MD). HCP funding was provided by the National Institute of Dental and Craniofacial Research (NIDCR), the National Institute of Mental Health (NIMH), and the National Institute of Neurological Disorders and Stroke (NINDS). HCP data are disseminated by the Laboratory of Neuro Imaging at the University of Southern California; (2) the *High-quality diffusion-weighted imaging of Parkinson’s disease* data base, Cyclotron Research Centre, University of Liège; (3) Hospital Gregorio Marañón (Madrid); (4) Cardiff University Brain Research Imaging Centre, provided by Chantal Tax.

## A. Noise Analysis of the Diffusion Measures

For the sake of simplicity in the analysis, we will estimate the bias of the cos^2^ in eq. (19). Note that it can be defined as

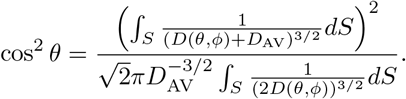

In this work we have approximated the surface integral by SH. However, for a noise analysis, it would be more convenient to use a summation, similar to what was done in Aja-Fernández et al. (2018):

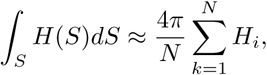

where *H*_*i*_ are N samples of H(S) uniformly distributed over the surface of the sphere. THis way, ve can write:

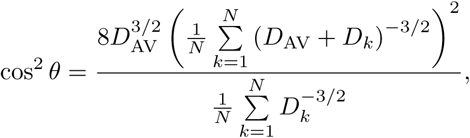

where *D*_*k*_ are samples of the signal *D*(*θ, φ*). We will assume that the SNR is high enough in the baseline, so we can consider it noiseless. We will assume that acquired signal *S*_*k*_ is corrupted by Rician noise, but, in the high SNR regime, it becomes a Gaussian. Thus, we can approximate the adquired signal as

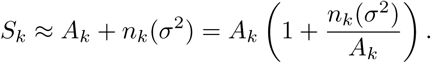

*A*_*k*_ is the original signal if no noise is present and *n*_*k*_(*σ*^2^) is the Gaussian noise, with zero mean and variance *σ*^2^. Under this assumption, we can write *D*_*k*_ as

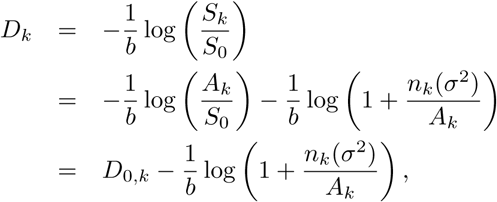

where *D*_0,*k*_ is the original diffusivity if no noise is present.

Our aim is now to calculate 𝔼{cos^2^ *θ*}. Let us define two variables:

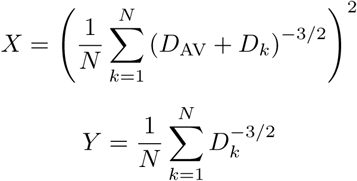

so that

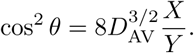

The expectation of cos^2^ *θ* can be written as Papoulis (1991)

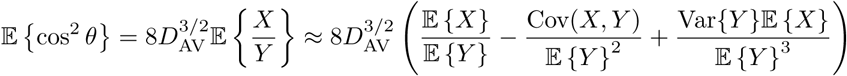

If we assume a high SNR we can make a first orden simplification. In order to calculate the bias, we will use a series expansion of *D*_*k*_, so that 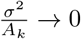:

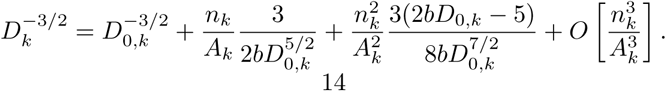

Similarly, we can also make the expansions for (*D*_*k*_ + *D*_AV_)^−3/2^ and (*D*_*k*_ + *D*_AV_)^−3^. After some algebra we can write:

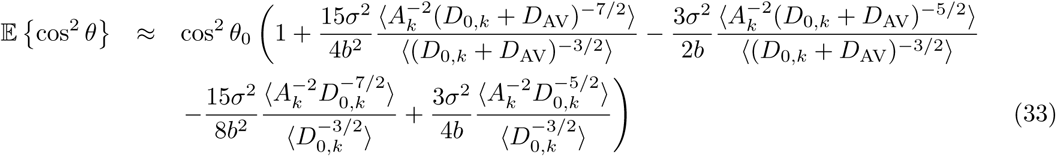

where cos^2^ *θ*_0_ is the original value (without noise) and 〈.〉 is the averaging operator:

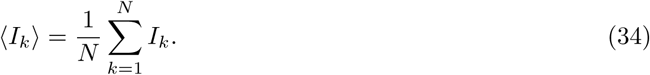

To better understand the effect of the SNR and the b-value, let us simplify eq. (33) by assuming an isotropic diffusion. In that case, *D*_0,*k*_ = *D*_AV_ for all *k*, and we can rewrite the expectation as:

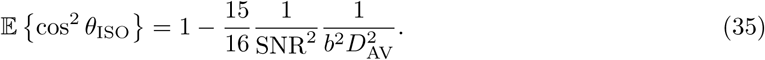

Therefore, the estimation bias of the cos^2^ *θ*_ISO_ can be quantified, for the isotropic case as:

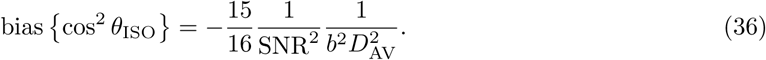

## Footnotes

2 The SNR of each of the individual baseline is high enough to make a Gaussian approximation feasible with a small error. Under this approximation we can assure that the average operator provides an unbiased output image (Aja-Fernández and Vegas-Sánchez-Ferrero, 2016).

3 https://www.nitrc.org/frs/?group_id=835.

4 mrtrix.org

5 The PA calculation is not available in the public distribution of DIPY. The current implementation has been kindly provided by Dr. Fick.

## References

Aja-Fernández, S., Pieciak, T., Tristán-Vega, A., Vegas-Sánchez-Ferrero, G., Molina, V., de Luis-García, R., 2018. Scalar diffusion-MRI measures invariant to acquisition parameters: a first step towards imaging biomarkers. Magn. Reson. Imag. In press.

Aja-Fernández, S., Vegas-Sánchez-Ferrero, G., 2016. Statistical Analysis of Noise in MRI. Springer.

Atkinson-Clement, C., Pinto, S., Eusebio, A., Coulon, O., 2017. Diffusion tensor imaging in Parkinson’s disease: Review and meta-analysis. Neuroimage: Clinical 16, 98–110.

Avram, A. V., Sarlls, J. E., Barnett, A. S., Özarslan, E., Thomas, C., Irfanoglu, M. O., Hutchinson, E., Pierpaoli, C., Basser, P. J., 2016. Clinical feasibility of using mean apparent propagator (MAP) MRI to characterize brain tissue microstructure. NeuroImage 127, 422–434.

Barrio-Arranz, G., de Luis-García, R., Tristán-Vega, A., Martín-Fernández, M., Aja-Fernández, S., 2015. Impact of MR acquisition parameters on DTI scalar indexes: a tractography based approach. PloS one 10 (10), e0137905.

Basser, P., Pierpaoli, C., 1996. Microstructural features measured using diffusion tensor imaging. J Magn Reson B 111 (3), 209–219.

Basser, P. J., 2002. Relationships between diffusion tensor and q-space MRI. Magnetic Resonance in Medicine 47 (2), 392–397.

Bernstein, A. S., 2019. Advanced diffusion mri techniques: Methodological development and clinical application. Ph.D. thesis, The University of Arizona.

Callaghan, P., Eccles, C., Xia, Y., 1988. Nmr microscopy of dynamic displacements: k-space and q-space imaging. Journal of Physics E: Scientific Instruments 21 (8), 820.

Descoteaux, M., Angelino, E., Fitzgibbons, S., Deriche, R., 2006. Apparent Diffusion Profile estimation from High Angular Resolution Diffusion Images: estimation and applications. Magn. Reson. Med. 56 (2), 395–410.

Descoteaux, M., Deriche, R., Le Bihan, D., Mangin, J.-F., Poupon, C., 2009. Diffusion propagator imaging: using Laplace’s equation and multiple shell acquisitions to reconstruct the diffusion propagator. In: International Conference on Information Processing in Medical Imaging. Springer, pp. 1–13.

Descoteaux, M., Deriche, R., Le Bihan, D., Mangin, J.-F., Poupon, C., 2011. Multiple q-shell diffusion propagator imaging. Medical image analysis 15 (4), 603–621.

Fick, R. H., Daianu, M., Pizzolato, M., Wassermann, D., Jacobs, R. E., Thompson, P. M., Town, T., Deriche, R., 2016a. Comparison of biomarkers in transgenic alzheimer rats using multi-shell diffusion MRI. In: International Conference on Medical Image Computing and Computer-Assisted Intervention. Springer, pp. 187–199.

Fick, R. H., Wassermann, D., Caruyer, E., Deriche, R., 2016b. MAPL: Tissue microstructure estimation using Laplacian-regularized MAP-MRI and its application to HCP data. NeuroImage 134, 365–385.

Gallager, R. G., 2008. Principles of digital communication. Cambridge University Press Cambridge, Cambridge,UK.

Hansen, B., Jespersen, S. N., 2016. Kurtosis fractional anisotropy, its contrast and estimation by proxy. Scientific reports 6, 23999.

Hosseinbor, A. P., Chung, M. K., Wu, Y.-C., Alexander, A. L., 2013. Bessel fourier orientation reconstruction (BFOR): An analytical diffusion propagator reconstruction for hybrid diffusion imaging and computation of q-space indices. NeuroImage 64, 650–670.

Hosseinbor, A. P., Chung, M. K., Wu, Y.-C., Fleming, J. O., Field, A. S., Alexander, A. L., 2012. Extracting quantitative measures from EAP: A small clinical study using BFOR. In: Med Image Comput Comput Assist Interv. Vol. 7511. Springer, pp. 280–287.

Mori, S., Wakana, S., Van Zijl, P. C., Nagae-Poetscher, L., 2005. MRI atlas of human white matter. Elsevier.

Ning, L., Westin, C.-F., Rathi, Y., 2015. Estimating diffusion propagator and its moments using directional radial basis functions. IEEE Trans Med Imag 34 (10), 2058–2078.

Özarslan, E., Koay, C. G., Shepherd, T. M., Komlosh, M. E., Irfanoğlu, M. O., Pierpaoli, C., Basser, P. J., 2013. Mean apparent propagator (MAP) MRI: a novel diffusion imaging method for mapping tissue microstructure. NeuroImage 78, 16–32.

Özarslan, E., Sepherd, T. M., Vemuri, B. C., Blackband, S. J., Mareci, T. H., 2006. Resolution of complex tissue microarchitecture using the Diffusion Orientation Transform (DOT). NeuroImage 31, 1086–1103.

Özarslan, E., Vemuri, B. C., Mareci, T. H., 2005. Generalized scalar measures for diffusion MRI using trace, variance, and entropy. Magn Reson Med 53 (4), 866–876.

Papoulis, A., 1991. Probability, random variables, and stochastic processes, 3rd Edition. McGraw-Hill.

Smith, S., Jenkinson, M., Johansen-Berg, H., et al., 2006. Tract-based spatial statistics: Voxelwise analysis of multi-subject diffusion data. Neuroimage 31, 1487–1505.

Tristán-Vega, A., 2009. A novel framework for the study of neural architectures in the human brain with diffusion MRI. Ph.D. thesis, Universidad de Valladolid, Valladolid, Spain, https://www.lpi.tel.uva.es/thesis.

Tuch, D. S., Reese, T. G., Wiegell, M. R., Wedeen, V. J., 2003. Diffusion MRI of complex neural architecture. Neuron 40, 885–895.

Wedeen, V. J., Hagmann, P., Tseng, W.-Y. I., Reese, T. G., Weisskoff, R. M., 2005. Mapping complex tissue architecture with diffusion spectrum magnetic resonance imaging. Magnetic resonance in medicine 54 (6), 1377–1386.

Westin, C.-F., Maier, S. E., Mamata, H., Nabavi, A., Jolesz, F. A., Kikinis, R., 2002. Processing and visualization for diffusion tensor mri. Medical image analysis 6 (2), 93–108.

Wu, Y.-C., Alexander, A. L., 2007. Hybrid diffusion imaging. NeuroImage 36 (3), 617–629.

Wu, Y.-C., Field, A. S., Alexander, A. L., 2008. Computation of diffusion function measures in q-space using magnetic resonance hybrid diffusion imaging. IEEE transactions on medical imaging 27 (6), 858–865.

Ziegler, E., Rouillard, M., André, E., Coolen, T., Stender, J., Balteau, E., Phillips, C., Garraux, G., 2014. Mapping track density changes in nigrostriatal and extranigral pathways in Parkinson’s disease. Neuroimage 99, 498–508.

